# *GWAIS-Web*: A Fast and Secure Web Service for Epistasis Detection in Genome-wide Association Interaction Studies

**DOI:** 10.1101/2023.05.16.540964

**Authors:** Lars Wienbrandt, Christoph Prieß, David Ellinghaus

## Abstract

**Motivation:** Genome-wide association interaction studies (GWAIS) are becoming increasingly important as estimates of genetic interactions at the genome-wide level using genome-wide data from hundreds of thousands of individuals from large biobanks showed that non-additive genetic variance plays a role in complex human traits in addition to additive genetic effects identified in genome-wide association studies (GWAS). However, a comprehensive genome-wide search for all combinations of second-order (SNPxSNP) or third-order (SNPxSNPxSNP) associations using millions of SNP markers is a very computationally intensive task, especially when hundreds of thousands or, in the near future, even millions of individuals can be studied with GWAS datasets. The runtime so far exceeds years, even if the search is performed on a multicore CPU server system.

**Results:** We developed *GWAIS-Web*, a web service for fast analysis of genome-wide interactions with case-control GWAS datasets. By using a hybrid combination of graphics-processing units (GPUs) and field-programmable gate arrays (FPGAs), *GWAIS-Web* speeds up epistasis detection methods for binary traits by a factor of more than 2000, allowing an exhaustive SNP-SNP GWAIS with a GWAS data set of one million SNPs and 500,000 individuals to complete within one day, which would take more than five years on a regular CPU server system. The user can choose between different methods for epistasis detection, such as logistic regression, BOOST, mutual information (MI) and others, with calculations in double precision and including on-the-fly filtering of correlated results based on linkage disequilibrium (LD). Due to the underlying common data structure of *GWAIS-Web*, all methods can be combined and processed together on-the-fly without increasing the runtime. The user can choose between 2nd order (pairwise) and 3rd order tests and can also limit the search to selected chromosomal regions. *GWAIS-Web* offers a high level of security through optional 2-factor authentication, encrypted connections and the protection of GWAS/user account data in accordance with the European General Data Protection Regulation (GDPR).

**Availability:** *GWAIS-Web* is freely available at https://hybridcomputing.ikmb.uni-kiel.de. The stand-alone software *HybridGWAIS* can be downloaded at https://github.com/ikmb/hybridgwais.

**Supplementary information:** Supplementary data are available online.

## Introduction

Genome-wide association studies (GWAS) have been very successful in identifying genotype-phenotype associations for more than a decade. However, it has been shown that single marker associations identified in GWAS explain only a fraction of the heritability of complex traits and diseases (Tam et al.; 2019) and do not reflect a realistic biological model, as single variants are tested individually for their association with the phenotype of interest, but not combinations of variants, although we assume that genetic associations of complex diseases or traits (and of potential comorbidities) belong to shared pathways with physically interacting biomolecules (Hu et al.; 2016), which is also reflected in the epistasis principle (Wei et al.; 2014). Statistical interaction refers to the interaction variance that can be explained by a combination of variants at different genetic loci (non-additive effects) rather than their independent single marker effects. For example, in psoriasis a significant interaction as measured by logistic regression has been detected between the genes *ERAP1* (rs27524) and *HLA-C* (rs10484554) with genotypic odds ratios (ORs) greater than 7.5 rel ative to the most protective two locus genotype (Genetic Analysis of Psoriasis Consortium et al.; 2010). The biological consequence of this interaction is that the *ERAP1* risk allele of rs27524 only has a strong effect in individuals carrying at least one copy of the risk allele of rs10484554 at the *HLA-C* locus. However, an exhaustive search for pairwise interactions between single nucleotide polymorphisms (SNPs) at the genome-wide level is computationally very time-consuming due to its runtime complexity *O*(*m*^2^*n*) (independent of the epistasis detection method used) for *m* number of variants and *n* number of individuals to be tested. With few exceptions where epistasis was successfully studied for variants pre-selected from established GWAS risk variants (The Australo-Anglo-American Spondyloarthritis Consortium (TASC) et al.; 2011; Kirino et al.; 2013; Keaton et al.; 2016), genome-wide association *interaction* studies (GWAIS) have not been as successful as GWAS because larger sample-sizes are also required to have sufficient statistical power on the genome-wide scale (Wei et al.; 2014). Nevertheless, this situation has changed since GWAS studies with hundreds of thousands or even millions of individuals have become possible (Evangelou et al.; 2018) and soon it will be possible to analyze over one million completely sequenced genomes (Saunders et al.; 2019). Studies using UK Biobank GWAS data already showed that a substantial portion of the unexplained heritability can be explained by non-additive interaction effects (Hivert et al.; 2021). All this opens up a huge new potential for conducting GWAIS, so we need to keep up with technical developments to make large-scale GWAIS computationally feasible.

To enable future large-scale GWAIS studies of the sample size of UK Biobank and beyond (i.e. for millions of variants and individual samples) to be carried out very precisely and with short runtimes, we previously combined algorithmic improvements of PLINK’s (gold standard) multiplicative logistic regression SNPxSNP interaction test (Purcell et al.; 2007; Chang et al.; 2015) with hardware acceleration and achieved a fundamental speedup (Wienbrandt et al.; 2019). Among other things, we could replace the formerly used covariate matrices in PLINK 1.9 with contingency tables and implemented the generation of these contingency tables on either a Field-Programmable Gate Array (FPGA) or a Graphics Processing Unit (GPU) with the calculation of the test statistic also on a GPU. One disadvantage, however, is that not all users have been able to benefit from the acceleration so far, because many users do not have access to these special high-performance computing resources to run our tool on FPGAs and/or GPUs.

We now introduce *GWAIS-Web*, a fast, convenient, secure and privacy-aware web service with a seamless integration of hardware accelerators that enables the genetics community to easily perform GWAIS for their genome-wide case-control studies without own compute resources. The web service is easy to use, has a convenient and secure user account system using only email address and access password, a job submission system, and sends email notifications about the completion of results. In addition, the web service has various security measures for the transmission and temporary storage of (sensitive) genetic data, including the exclusive use of encrypted and certified connections, password-protected data access, optional 2-factor authentication (2FA) and the protection and self-deletion of all user account data in accordance to the European General Data Protection Regulation (GDPR). Besides PLINK’s logistic regression test, which we have now also implemented for SNPxSNPxSNP (i.e. third-order) interactions in *GWAIS-Web* with runtime complexity *O*(*m*^3^*n*), we have added several other second- and third-order epistasis screening methods and features to *GWAIS-Web*, for example, the ability to run multiple interaction tests simultaneously in one run (without increasing the runtime) and region selection options to focus on chromosomal regions of interest (e.g. interaction testing of SNPxSNP or SNPxSNPxSNP tuples only within known GWAS risk loci). We are also able to distribute the computational load to GPU and FPGA accelerators in an automated way, which drastically reduces the runtime. We make all new implementations and GPU and FPGA hardware resources freely available in *GWAIS-Web* at https://hybridcomputing.ikmb.uni-kiel.de. The stand-alone software *HybridGWAIS* (with or without GPU/FPGA support) can be downloaded at https://github.com/ikmb/hybridgwais.

## Materials and methods

### *GWAIS-Web* server infrastructure

An overview of the server infrastructure is illustrated in **Figure 1**. The server infrastructure of *GWAIS-Web* is divided into two main components referred to as *frontend* and *backend* system. The frontend hosts the web service, a database with user information, and the filesystem for storing uploaded files and analysis results. It is directly connected via a dedicated network connection to the backend system which provides a job queuing system and the necessary computing power supported by FPGA and GPU hardware accelerators for processing the user’s submitted jobs. The division into frontend and backend system allows job processing to be independent from web service operations and keeps the web service functional despite a full computational load on the backend during job processing. For more technical details, see **Supplementary 1.1**.

**Figure 1.**
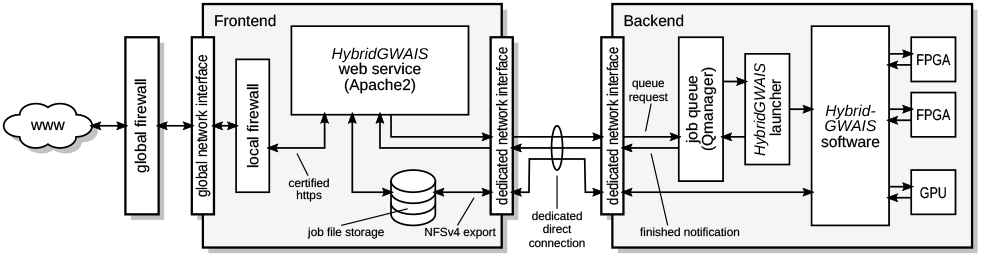
The *GWAIS-Web* server architecture consists of two main components: the frontend and backend systems. The web service is hosted by the frontend while jobs are processed on the backend system. The processing order is controlled by the job queue which we implemented in the *Qmanager* software on the backend. Files are stored on the frontend and accessed via an *NFSv4* network file system. **Supplementary 1.1** presents technical details of the setup of the frontend and backend systems.

### Web user interface

To use our web service, an account registration with not more than a valid email address and a password for authentication is necessary. The account is required to manage job submissions, to send email notifications (e.g. when a job is finished), and to provide the result files to the user for download. To enhance security, it is possible to register a key device for 2-factor authentication. A typical GWAIS job submission is started by uploading case-control genotype data in PLINK’s .bed/.bim/.fam format first, which can be done via the web browser, a provided URL or via SFTP. Then the user can choose from different epistatis detection methods (either for 2nd-order (pairwise) or 3rd-order interactions) including logistic regression, BOOST (Wan et al.; 2010), log-linear test and entropy-based measures such as mutual information and information gain. Further, the linkage disequilibrium (*r*^2^) measure can optionally be added to the selected methods for on-the-fly calculation and/or on-the-fly filtering of results, as well as many other runtime options. The job progress can be supervised by the user when logged-in, and the job continues to run even after logout. When the run is finished, the service sends a notification email to the user that the results are available for download from the user’s job administration section. The download can be started either directly in the browser or via a script in a command line terminal. This workflow is illustrated in **Figure 2**.

**Figure 2.**
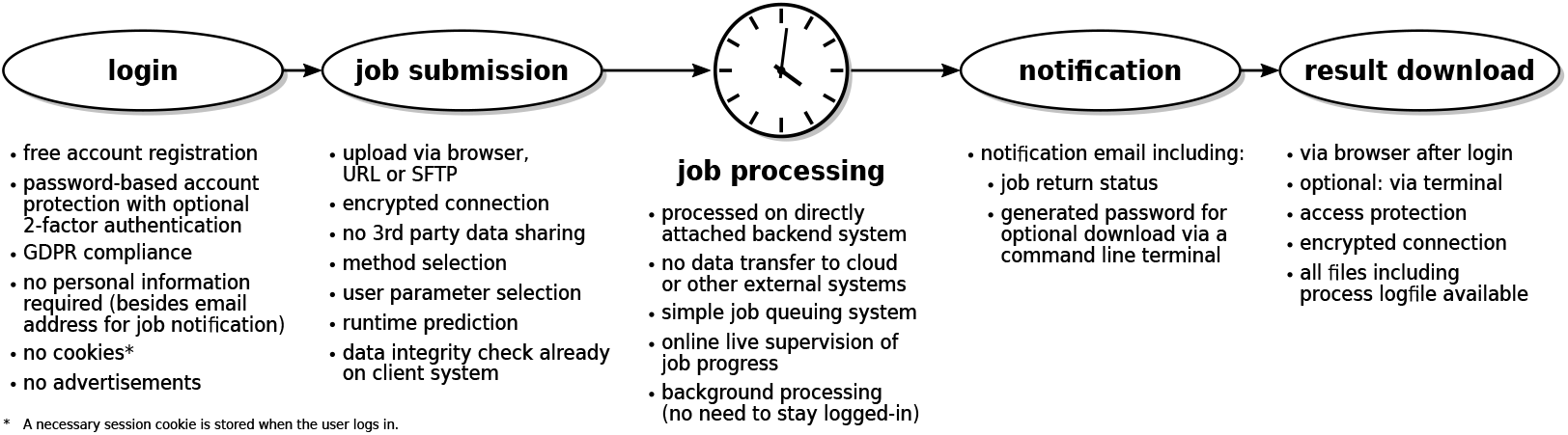
Epistasis screening workflow in *GWAIS-Web*. After registering an account and logging in, the user uploads their dataset and completes the simple job submission form by selecting the desired test methods, chromosomal regions (optional) and algorithmic options. The job is then queued and processed on the directly connected backend compute system. Once the calculation is complete, the user receives a notification email informing them of the return status. Result downloads are possible either via the browser or a command line terminal.

For details on the user interface, its functionality and implementation, please refer to **Supplementary 1.2**.

### Protection of personal data

Because genome-wide data is personal data, we must ensure the requirements of privacy by design (Pardau and Edwards; 2017). There are many international examples of legislation implementing privacy by design. In the US, this principle has been enshrined in the California Consumer Privacy Act (CCPA), among others, and in 2012 the US Federal Trade Commission (FTC) published a framework of best privacy practices for implementing privacy and data security for businesses that collect and use consumer data (Federal Trade Commission; 2012). In Europe, privacy by design is explicitly mandated in the European Union’s General Data Protection Regulation 2016/679 (GDPR) (The European Parliament and the Council of the European Union; 2016). How we comply with the rules of the GDPR is explained in detail on our web services website. In summary, we do not use the provided email address for other purposes than job and account management, we do not use the uploaded genetic data for other purposes than the execution of the analysis request, and no one than the user has access to the data. All data remains locally on our university server located in Kiel, Germany, and all uploaded files and result files (except log and status files) are automatically deleted 7 days after completion of the corresponding job. No data is passed on to third-party services, the website is free of advertising and does not store any cookies on the user’s computer (except for a technically necessary session cookie). For reasons of transparency, the user can also download all personal data stored on our server in a single file with a single mouse click. All user-related data is immediately deleted if the user decides to delete their account (which can be done directly by the user in the user administration section).

### The *HybridGWAIS* software and enhancements

The *HybridGWAIS* software used in *GWAIS-Web* is freely available as a stand-alone software at GitHub: https://github.com/ikmb/hybridgwais. The core idea of *HybridGWAIS* was to accelerate PLINK’s (Purcell et al.; 2007; Chang et al.; 2015) multiplicative logistic regression epistasis detection test by reducing the computational effort by using a contingency table instead of a covariate matrix (Wienbrandt et al.; 2019). By creating the contingency tables on an FPGA accelerator and running the statistical tests on a GPU, we had achieved a speedup of a factor of more than 1000x compared to the original PLINK 1.9 software, which we have now even further improved to achieve a speedup of more than 3000x (in the order of 50,000 variants and 100,000 samples). To support GWAIS researchers who do not have access access to FPGA accelerator hardware or do not want to use *GWAIS-Web*, we have implemented *HybridGWAIS* that it can optionally be used with GPU accelerators alone (then reaching a speedup of more than 300x for the above example) or parallelized across multiple threads to be used without hardware accelerators on a desktop PC or computing cluster node with a multicore CPU (with a 10x speedup).

Since case-control datasets can generally have genotype information condensed into contingency tables, many different kinds of statistical epistasis tests can be conducted on them. In *HybridGWAIS*, we therefore re-use the generated contingency tables for all user-selected epistasis tests, and as the contingency table generation takes most of the computational time, the analysis of a dataset can be done by conducting several tests at the same time without significant increase in runtime. Besides PLINK’s multiplicative logistic regression test, we implemented BOOST (Wan et al.; 2010), log-linear regression (i.e. BOOST without pre-filtering), and the entropy-based tests mutual information and information gain. Because ideally the variants to be tested should not be in linkage disequilibrium (LD) with each other, an LD-test calculating the *r*^2^-value between two markers can optionally be added in addition to the selected methods. The user may also select any *r*^2^-threshold to filter variant pairs in LD on-the-fly (though only the specific pair in high LD will be filtered out; the variants of this pair will still be used for pairings with other variants and thus remain available for the exhaustive search). We also included third-order tests to analyze marker triples instead of only marker pairs. The LD-test and LD-filter then apply to all pairwise combinations in each marker triple. The results of all tests are summarized in a table showing the *n* best results according to the test selected first. (The parameter *n* is freely selectable by the user.) The result table is implemented with the help of a *MinMaxHeap* where the minimum and maximum elements can be accessed in constant time and the insertion of an element requires logarithmic (*O*(log *n*)) time.

Furthermore, *HybridGWAIS* allows for arbitrary selection of chromosomal regions (e.g. only known GWAS regions) to conduct only a subset of all possible tests, thus significantly reducing the analysis runtime and the search space. Regions can be selected independently for each pair/triple index, making it is possible to test only markers from a certain chromosome or genetic region (e.g. from the human leukocyte antigen region (HLA)) to another chromosome or genetic region. Each region can be specified as an interval with any start and end position in the input dataset. It is also possible to additionally exclude a region with arbitrary start and end position, either completely or only for the first marker. This allows analyses e.g. for a selected chromosome against the rest of the dataset (excluding the chromosome) or analyses of one specific region against another region (overlapping or not). Users can set a *proximity exclude range* measured in base pairs. If two markers of a pair or triple are within this range, the combination will not be selected for testing. This can be helpful when marker pairs that are close to each other generate a high statistical score but are unlikely to form an epistatic relationship due to their proximity.

With the ability to focus only on specific regions, *HybridGWAIS* is also able to automatically distribute the analysis across multiple FPGAs to further increase computational speed. The web service takes advantage of this by using up to two FPGAs for one analysis, depending on the size of the GWAS dataset. Thus, there is no variant limit induced by the FPGA memory, as a large GWAS dataset is distributed over several chunks before being processed by an FPGA.

We present a list of our newly implemented features in comparison to PLINK 1.9 in **Table 1**. Further details on the implementation and acceleration of *HybridGWAIS* can be found in **Supplementary 2** and **3**.

**Table 1.**
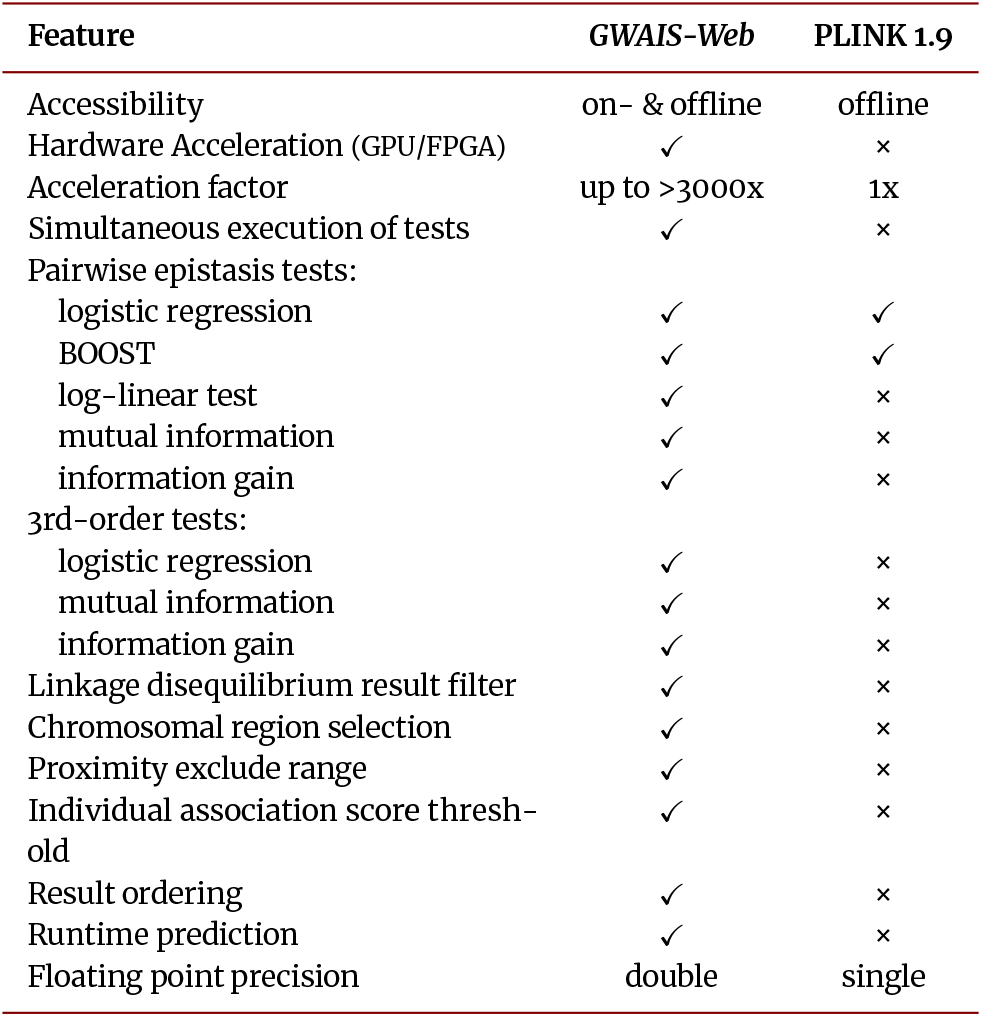
Feature comparison of *GWAIS-Web* versus PLINK 1.9.

### Benchmark Settings

For runtime benchmarking, we used simulated datasets of different sizes with a varying number of markers and samples, but with an equal distribution of cases and controls (see **Supplementary 4.1** for details on how the benchmark data were created). It is important to remember that the most computationally intensive part is the creation of the contingency tables, which is independent of the content of the table. Thus, the total runtime essentially depends only on the size of the dataset, not on its contents or the type of statistical test. (In particular, the content of the tables have a minimally negligible impact on the runtime, since in logistic regression, for example, the number of Newton iterations depends on the quality of the result, but is internally capped by a number of 16 iterations anyway).

Consequently, we initially applied the benchmark process only with the logistic regression method to different combinations of sample and marker sizes. We tested the *HybridGWAIS* software with different accelerator methods on our backend system, i.e. (a) using only multi-processing with 32 threads on our CPUs alone, (b) using one GPU accelerator, (c) using a combination of one FPGA and one GPU accelerator, and (d) using two FPGA accelerators and one GPU accelerator in combination. (Note that the web service uses only the fastest configuration (d) for regular operations.) We also performed PLINK analysis on the smaller datasets to measure the speedup of *HybridGWAIS* over PLINK 1.9 (Chang et al.; 2015). (Note that in the newer PLINK 2.0 software, epistasis testing is not yet implemented.)

To illustrate that the combination of different test methods has a negligible impact on the runtime, we ran several tests with the same dataset (100,000 samples with 100,000 variants), but different combinations of test methods. We ran this benchmark with the same accelerator configurations (b)-(d) as above. Here, we omitted the CPU-only configuration (a) because the expected runtime was too high and we did not expect different results in terms of runtime deviation compared to the configurations with accelerator(s) used. We measured all runtimes with the standard GNU time command.

## Results and discussion

**Figure 3** shows the runtimes and speedups compared to PLINK 1.9 for the pairwise logistic regression test of our benchmarks, firstly for a fixed number of *n*=50,000 samples and a variable number of variants. Secondly, we demonstrated by linear extrapolation how the benchmark would behave for a fixed number of *n*=500,000 samples. In **Figure 4** we additionally illustrated the benchmark for a fixed number of *m*=50,000 variants and a variable number of samples. The exact runtime measurements of our benchmarks can be found in **Supplementary 4.2**. As expected, the runtime of *HybridGWAIS* increases quadratically with the number of variants and linearly with the number of samples. PLINK suprisingly shows a more than linear runtime increase with increasing sample size, at least for the sample size we used in our benchmarks, which was less than 100,000 (since PLINK’s runtime increased impracticably for a larger sample size). This observation is underlined by the increase in speedup with an increasing number of samples.

**Figure 3.**
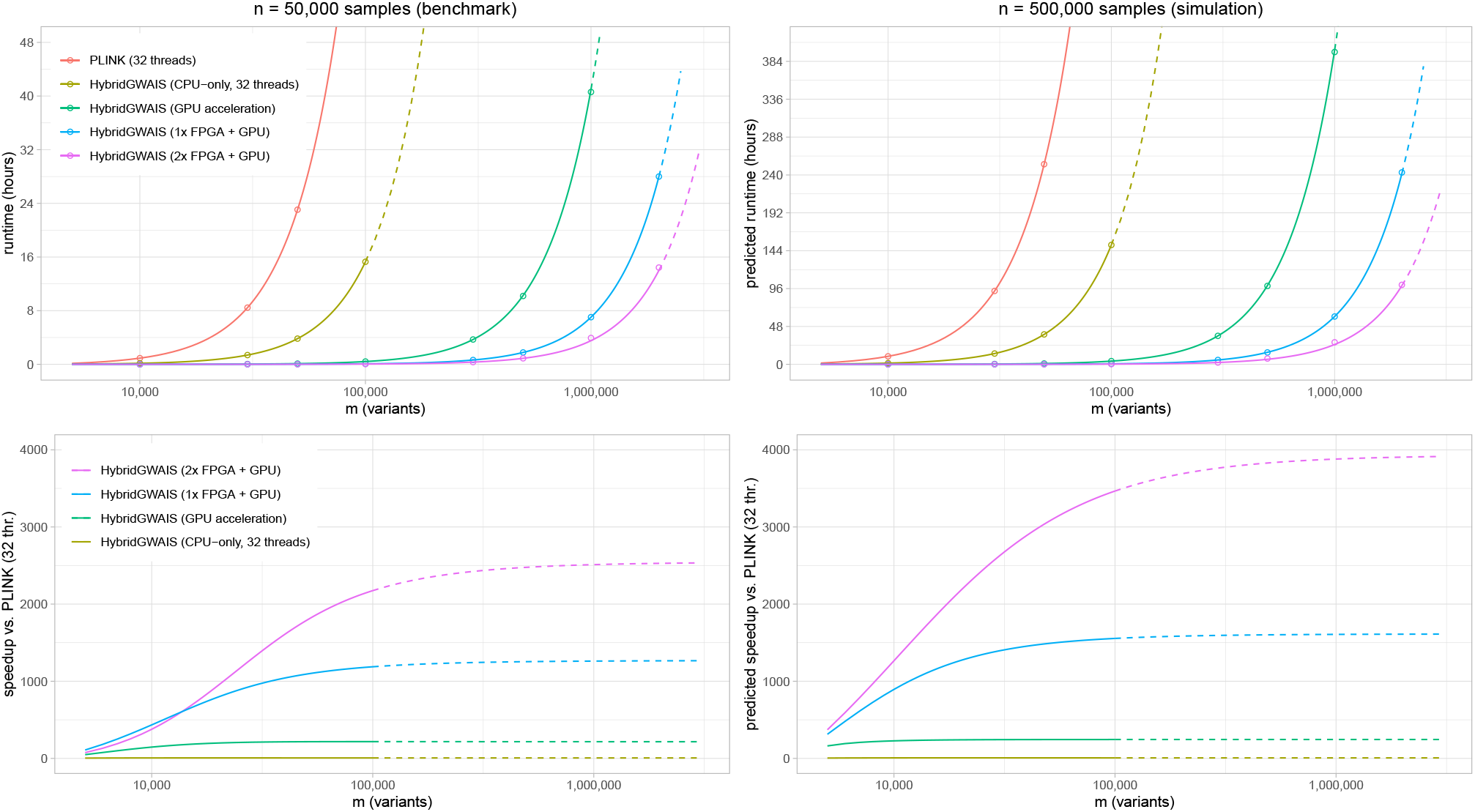
Benchmark and simulation results from *HybridGWAIS* compared to the PLINK epistasis test, using 2nd-order logistic regression interaction testing as an example. The upper left subfigure shows the runtimes of a fixed number of *m*=50,000 samples and a variable number of variants of the epistasis test in PLINK 1.9 running with 32 threads and four different acceleration modes in *HybridGWAIS*: (a) CPU-only multi-processing with 32 threads, (b) acceleration with one GPU, (c) combined acceleration with one FPGA and one GPU, (d) combined acceleration with two FPGAs and one GPU. The lower left subfigure shows the respective speedups of the four modes in *HybridGWAIS* compared to PLINK 1.9 with 32 threads. The right subfigures show a simulated benchmark by illustrating the predicted runtimes and speedups for a fixed number of *n*=500,000 samples and a variable number of variants. The x-axes of all subfigures are in logarithmic scale. Solid lines in the runtime graphs represent a quadratic fitting of the measured (upper left subfigure) and predicted (upper right subfigure) data points, dotted lines the extrapolation of these functions. Solid lines in speedup graphs represent the relation of PLINK’s fitted runtime function to the fitted functions of each acceleration mode respectively, dotted lines represent the relation of PLINK’s runtime extrapolation to the fitted functions of each acceleration mode.

**Figure 4.**
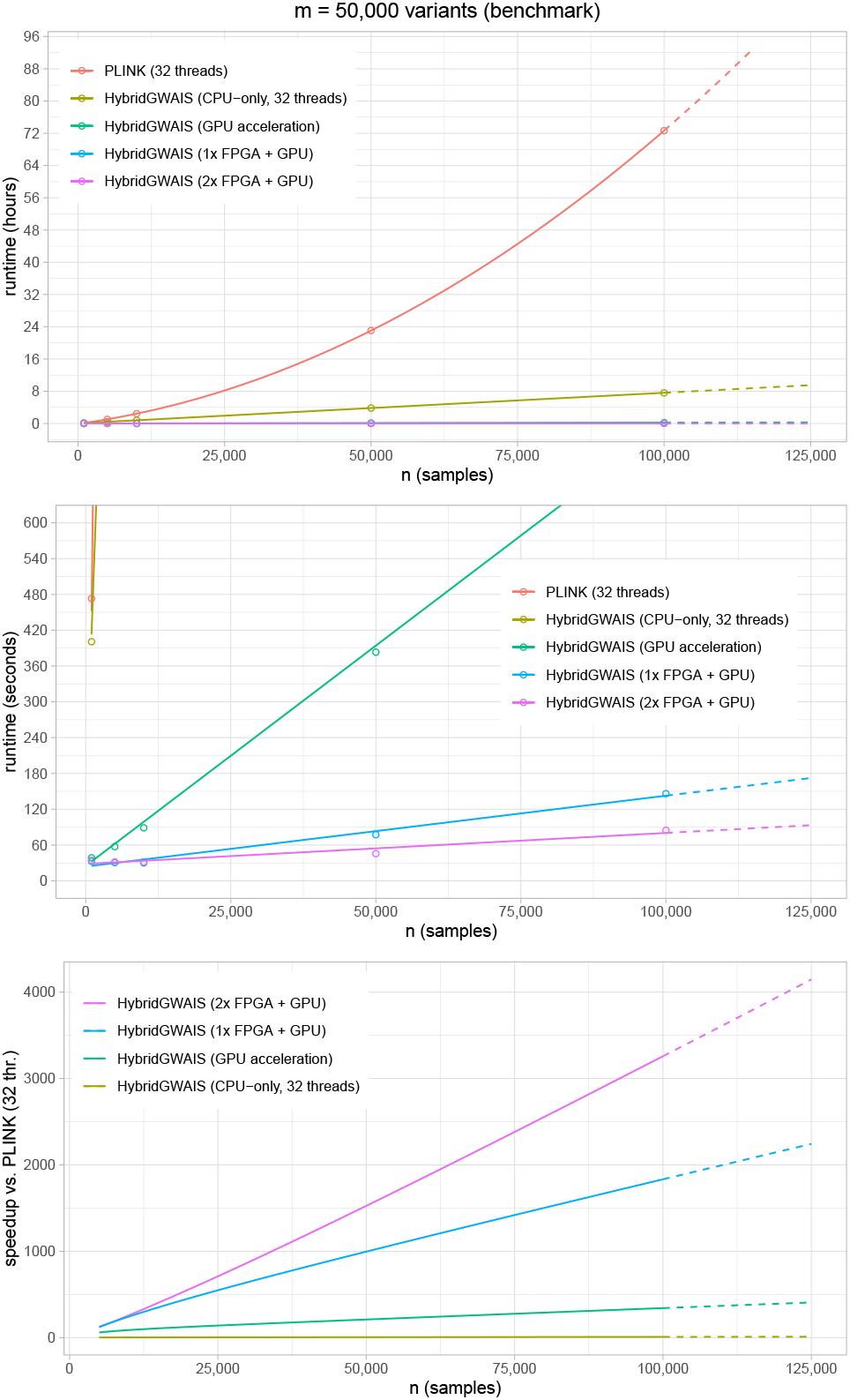
Benchmark results from *HybridGWAIS* compared to the PLINK epistasis test, using logistic regression as an example. The top figure shows the runtimes of a fixed number of *m*=50,000 variants and a variable number of samples of the epistasis test in PLINK 1.9 running with 32 threads and four different acceleration modes in *HybridGWAIS*: (a) CPU-only multi-processing with 32 threads, (b) GPU acceleration, (c) combined acceleration with one FPGA and a GPU, (d) combined acceleration with two FPGAs and a GPU. The second figure shows a scaled version with a magnified y-axis to highlight the differences between the last three acceleration modes. As a consequence, most PLINK and CPU-only data points are not in the visible region anymore. The bottom figure illustrates the corresponding speedups versus PLINK 1.9 (using 32 threads). Solid lines indicate the fitted interpolation functions between data points, dotted lines indicate an extrapolation.

In Figure 3, we observe a speedup of more than 2100x (exactly 2172x) for *n*=50,000 samples and *m*=100,000 variants when using two FPGAs plus GPU acceleration compared to PLINK 1.9. This speedup is predicted to reach about 3500x in our simulation for an increased number of *n*=500,000 samples and the same number of variants. Figure 4 shows that the measured speedup is already more than 3000x (exactly 3084x) for *n*=100,000 samples and *m*=50,000 variants. The speedup factors in this scenario for the other test configurations are 1790x for using one FPGA plus one GPU, 346x for GPU-only and 10x for CPU-only processing. In absolute runtimes, for this example we measured a runtime of 1 minute and 25 seconds for two FPGAs plus one GPU acceleration, 2 minutes and 26 seconds for one FPGA plus one GPU acceleration, and 12 minutes 36 seconds for GPU-only acceleration. The CPU-only runtime was 7 hours 37 minutes, resulting in 323x, 188x and 36x acceleration when comparing both FPGA+GPU accelerations and GPU-only acceleration against the CPU-only run. In contrast, the absolute PLINK 1.9 runtime for this example exceeds 3 days, highlighting the need for runtime acceleration.

It is worth noting that for datasets with a smaller number of variants, the observed speedup when using two FPGA accelerators is lower than the speedup when using only one FPGA accelerator (see Figure 3, fixed number of *n*=50,000 samples and variants below 17,000). We interpret this as a consequence of the overhead caused by distributing the analysis space over two FPGAs, which is not necessary when using only one FPGA.

To verify the correctness of our results, we confirmed that the order of our reported results matched the reported PLINK results (after sorting by score). The result scores and reported p-values of *HybridGWAIS* equal the results from PLINK’s logistic regression tests (apart from minor deviations resulting from the fact that in *HybridGWAIS* we use calculations in double precision floating point format, wherease the PLINK software only uses single precision format).

For the second benchmark (measuring the impact of different and multiple test methods on the runtime), the table of runtimes is shown in **Supplementary Table 7**. It can be clearly seen that the runtime in the accelerator configuration (d) ranges from 285 seconds (using e.g. mutual information tests alone) to 289 seconds (for the combination of all available methods) which means a deviation of only -0.43% to 0.9% compared to an analysis using the logistic regression method alone (286 seconds). Similar results are observed for the other accelerator configurations, showing that the selection of the test method actually has only a negligible influence on the runtime.

## Conclusion

With *GWAIS-Web*, we offer a fast and convenient web service for performing genome-wide interaction studies on large GWAS datasets. By using our cost-free web service, researchers can benefit from the superior performance of our FPGAs+GPU acceleration without having to purchase and install the necessary hardware themselves. Furthermore, we offer a variety of different epistasis test methods (currently to analyse binary traits) that can be run in parallel without a further runtime delay. The new features for selecting arbitrary chromosomal regions and for LD-filtering enable users to reduce the search space for interactions as well as to remove statistically questionable results from the result. This may also save valuable runtime compared to performing a complete genome-wide screening with a much larger result list and then filtering the results.

The convenient web interface allows users to easily run a genome-wide epistasis screening analysis without having to deal with software installation and command line parameters. With GDPR-compliance, data and account security, transparency, locality, no links to external websites, no cookies and other forms of data collection, researchers can quickly benefit from their analysis results without worrying about safety or being bothered by unwanted advertisements.

The web service will be continuously maintained and extended in future projects to include other statistical methods, for example, BiForce (Gyenesei et al.; 2012) and the joint-effects test (Ueki and Cordell; 2012). Currently, we are working on implementing linear regression for quantitative traits. Further, we plan to include permutation-based testing methods such as MB-MDR (Calle et al.; 2007) and corrections for covariates (such as principal components from principal component analysis (PCA)) as well as support for processing genotype dosage data from genotype imputation.

## Availability of source code and requirements

The *GWAIS-Web* web service, accessible at https://hybridcomputing.ikmb.uni-kiel.de, consists of the combination of three separate projects: (a) *HybridGWAIS*: standalone software for epistasis detection with optional GPU support; (b) *HybridGWAIS-FPGA*: The source code for the FPGA hardware design to be used in *HybridGWAIS*; (c) *Qmanager*: The source code for our job queueing system.

- Project name: HybridGWAIS
- Project home page: https://github.com/ikmb/hybridgwais
- Operating system: Linux
- Programming language: C++ (incl. CUDA), PHP, JavaScript, SQL, bash for *GWAIS-Web*
- Other requirements: BOOST, TBB, Cmake
- License: GNU GPL v3.0

The source code of the FPGA design is available at https://github.com/ikmb/hybridgwais-fpga. To compile the FPGA design the Xilinx Vivado development platform is recommended. Note that the required IP cores are not provided due to licensing restrictions and must be re-generated by the user.

The *Qmanager* software is available at https://github.com/ikmb/qmanager. To setup and run this software independent of our web service, a Linux operating system with packages available from standard distributions is required. See listed URL for details.

## Ethical approval

Not applicable.

## Data availability

All runtime measures from our benchmarks are listed in the **Supplementary Material**.

