## Supplementary Material for "*GWAIS-Web*: A Fast and Secure Web Service for Epistasis Detection in Genome-wide Association Interaction Studies"

May 16, 2023

### 1 Implementation of *GWAIS-Web*

#### 1.1 *GWAIS-Web* server infrastructure

An overview of the server architecture of *GWAIS-Web* is depicted in **Figure 1** in the main paper. It is divided into two components, referred to as *frontend* and *backend* system, to make our service fast, stable and secure. The frontend hosts the web service including a database with user information, such as login credentials and all data regarding submitted jobs. Uploaded files and result files are also stored on this system. The frontend is equipped with an Intel Xeon Silver 4110 8-core CPU @ 3 GHz and 128 GB RAM. It offers more than 7 TB of redundant storage capacity and is currently running on an Ubuntu 22.04 LTS Linux system, which is regularly updated. The web service is mainly written in *PHP* and *JavaScript* and is hosted by an *Apache2 v2.4* [1] server. The database is implemented in *PostgreSQL* [2], version 13.7.

With two Intel Xeon E5-2667 v4 8-core CPUs running at @ 3.6 GHz and 512 GB RAM, the *backend* provides the necessary computing power for the actual processing of the submitted jobs. The operating system is Ubuntu Linux 22.04 LTS as well. The backend is further equipped with two FPGA accelerators (Alpha Data ADM-PCIE-8K5 PCIe containing a Xilinx Kintex UltraScale KU115 FPGA each), and an Nvidia Tesla P100 GPU. The backend's main tasks are to host the job queue, that we have implemented in our simple but efficient job queueing system *Qmanager* [3], and to run the *HybridGWAIS* tool [4] to process the user's epistasis screening jobs. *HybridGWAIS* is written in *C++* using *CUDA* and compiled with *GCC v11.3.0*, the *Qmanager* is implemented in *Rust*.

To ensure direct communication without potential interception risks and routing problems, the frontend and backend are connected via a direct Ethernet connection and dedicated network devices. Communication between these systems via this connection is limited to two applications: First, the backend offers access to the *Qmanager* via a single open port on this direct connection to the frontend. Second, the frontend only exports the storage file system via *NFSv4* to the backend via this connection and accepts notification calls from the *Qmanager*. A firewall on the frontend (implemented with the Linux system tool *iptables*) rejects all incoming traffic from the internet except *https* requests. (In particular, *http* requests are also allowed, but are automatically redirected to *https*.) The backend firewall is configured to completely block all incoming traffic from the internet. Only administrative access via *SSH* is exceptionally allowed only from the local network for both systems.

Since the servers are located in the infrastructure of Kiel University, an additional firewall from the university's router ensures that rules are not violated. The server's certificate (required for secure *https* connections) is issued via the University by the external organization *GEANT Vereniging* and will be verified by any browser's standard certificate chain.

#### 1.2 User interface and functionality

##### 1.2.1 Registration and login

*GWAIS-Web* requires the registration of a personal user account including a valid email address. The user account enables the protection against unauthorized access by third-parties to user data and test results. The email address is used to inform the user about her/his job events (such as a job completion, because a genome-wide screening process may take several hours). We explicitly point out that we do not use the provided email address for purposes other than job notifications and account management and do not collect any usage information or statistics of our service in connection with user accounts.

To complete the registration process, the user gets a verification email from our server to the provided email address. (For sending mails the tool *mSMTP* is used.) The email contains a one-time link that finally validates the email address and activates the user account. It is valid for 7 days, after which the account will automatically be deleted if it was not activated before.

The account protection is implemented either via a simple password or optionally for extra security, the user may register a key device for 2-factor authentication (see **Sect. 1.2.5** below).

After login, the user is able to submit jobs, manage ongoing or completed jobs, download results, and to manage the account. Note that the web service will logout the user automatically after 30 minutes of inactivity.

In general, we renounce the usage of cookies, but to verify the current login status we need to store a necessary session cookie.

##### 1.2.2 Job submission

New jobs are arranged in a queue to ensure a fair order of execution among users (on a first-come, first-served basis). To run a job within our service, the upload of case-control genotype data in PLINK's `.bed/.bim/.fam` format is required. A screenshot of the job submission form is depicted in **Supplementary Figure 1**. Uploading a file triplet is possible in three different ways. The easiest way is to upload via a browser. Files can simply be selected in a file selection dialog or dragged and dropped in the designated box. Alternatively, the web server can actively fetch the files from public URLs (e.g. pointing to a private server) that are provided in the submission form. Or the upload via *Secure File Transfer Protocol (SFTP)* can be chosen, which requires the user to provide the host URL, login credentials and relative paths to the files on that server. Note that we use the login credentials only for the purpose of downloading the files. We never submit them in plain-text, as SFTP is an encrypted connection, and we delete them immediately after the upload to our server has finished.

During the upload process the files are checked for consistency to improve reliability and ensure stable job runs. In particular, in the case of a browser upload, a JavaScript module executed on the client's device checks selected files already before upload. The user gets an immediate feedback about files not matching the required restrictions:

- A file triple in PLINK's bed/bim/fam format must be uploaded with the correct endings `*.bed`, `*.bim` and `*.fam`.
- The maximum size of the `.bim`-file or the `.fam`-file must not exceed 100 MB.
- The `.bim`-file and the `.fam`-file are parsed on the client side to determine the number of variants, and the number of cases and controls beforehand. Thus, these files need to have at least 6 columns. Data that contains no cases or no controls is rejected beforehand. The same applies to files with no data at all.
- The `.bed`-file is checked for consistency, i.e. it must contain PLINK's file signature in the first three bytes and the file size must conform to the number of markers in the `.bim`-file and the number of samples in the `.fam` file.

If the user selects another upload method, the files are checked on our server while uploading and the upload stops immediately if a file is not valid.

The `.bim` file is also used to enable the selection of regions of interest within the provided markers (see Sect. 3.4). Other runtime options include an arbitrary job name, the number of results in the output list, and, certainly, the selection of epistasis testing methods for this job including a linkage disequilibrium ($r^2$) filter. (Currently, *GWAIS-Web* supports the methods listed in Table 1 in Sect. 2.2.)

When submitting the job, the values in the form are translated by a PHP script in the background to the command line options required for the *HybridGWAIS* software. The relative location of the job folder, that is uniquely created for the input, output and log files, is submitted along with the command-line options to the job queuing system *Qmanager* (see Sect. 1.2.3).

The queuing status and progress of the job can be supervised in the *Jobs* section. Once a job has finished, the web service gets notified via a secured notification URL (allowing only connections from the backend system). According to the jobs return state (success or failed), the web service generated download links for the result files and notifies the user per email (see Sect. 1.2.4 for details).

##### 1.2.3 Job queuing system

We implemented a simple and freely available job queuing software in the *Rust* programming language named *Qmanager* [3]. Its basic features are the reception of commands that should be processed in order on the backend system. It handles all queued, running and finished jobs (including their output and return codes) from the frontend system.

Job queuing request are handed over via a *https* connection in *JSON* format. For security reasons, the *Qmanager* is not allowed to run any arbitrary command. Accepted commands have

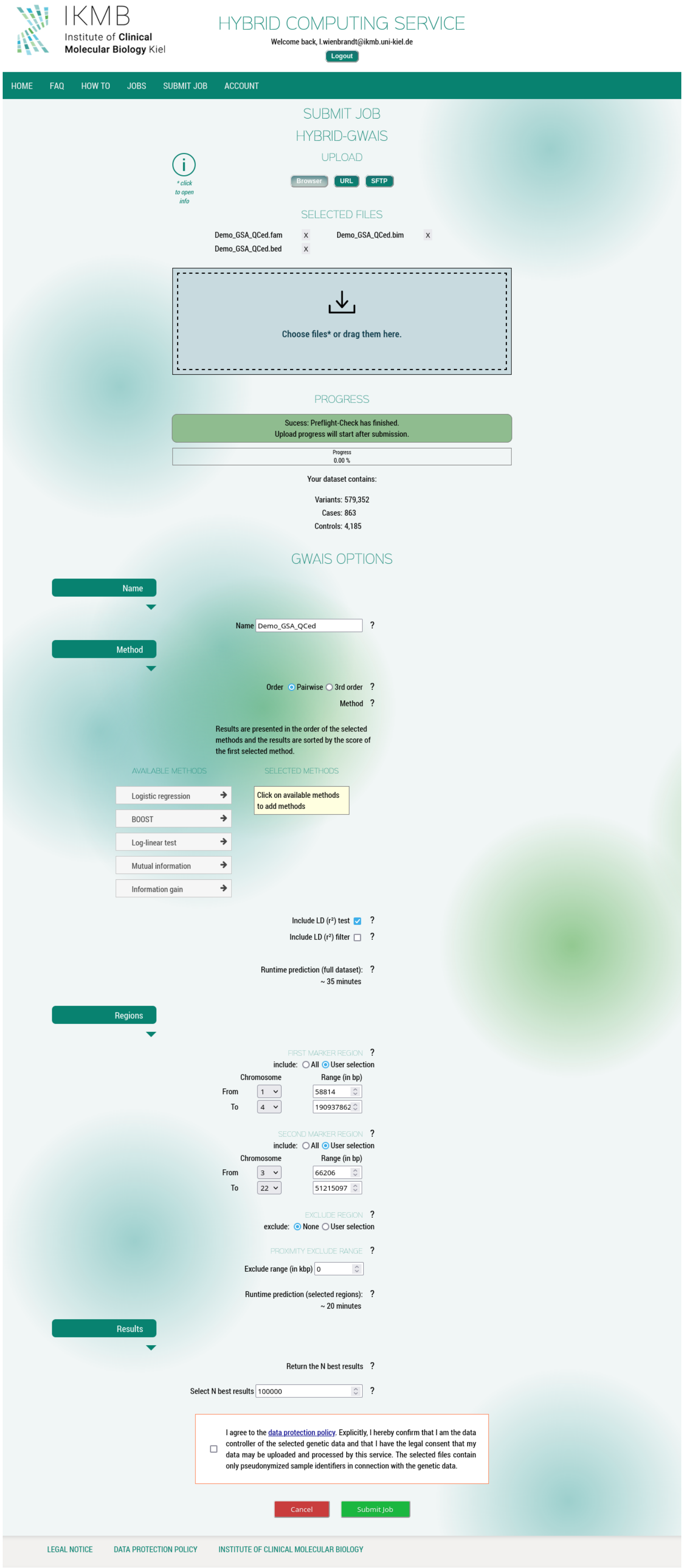

Supplementary Figure 1: Screenshot of *GWAIS-Web*'s job submission page with the available configuration and region selection options.

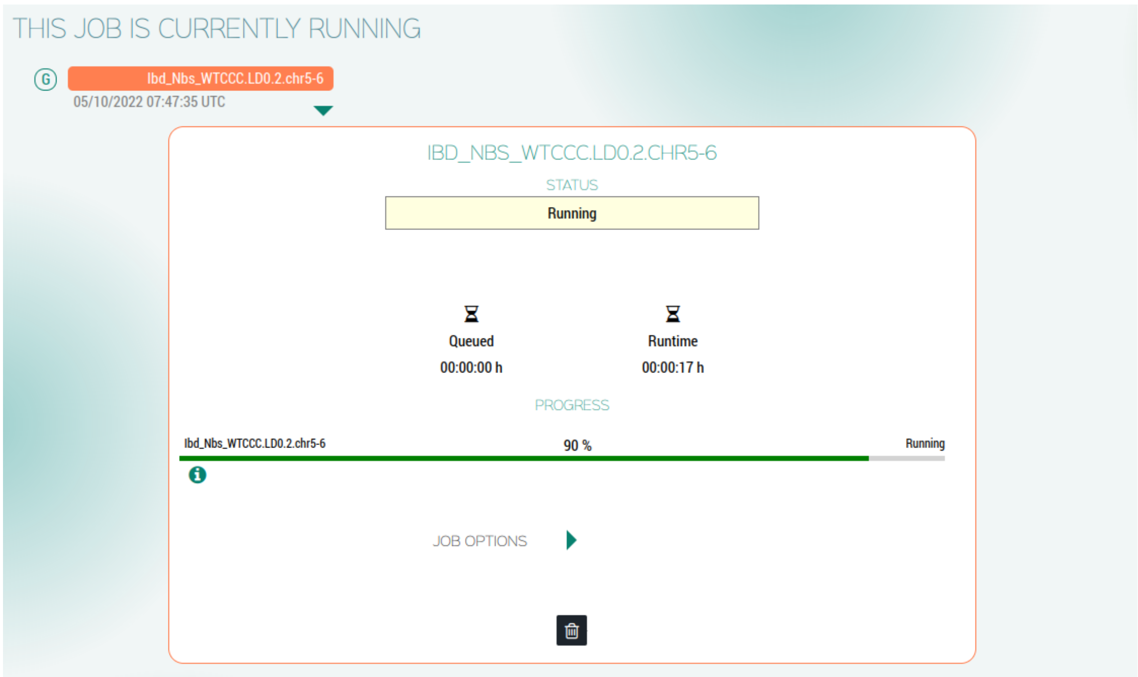

Supplementary Figure 2: Screenshot of an exemplary job progress in *GWAIS-Web*.

to be registered in the configuration file of the *Qmanager* and are accessed via an identifier, not the command itself. Parameters though are passed through directly.

For *GWAIS-Web* the only registered command is the call to a *HybridGWAIS* launch script. The script parameters are the (relative) location of the user's input data and the user options from the web interface. The script in turn converts the relative location to an absolute path and launches the *HybridGWAIS* executable for the user's input files with added default and user options.

The *Qmanager* is run as a system service and provides options to submit a job to the queue, to remove jobs from the queue (either enqueued or finished) and to kill a running job. Whenever a job terminates (finished or killed), the *Qmanager* calls a pre-configured notification URL with the job identifier. In this case, the notification script is executed on the frontend system and causes several operations such as changing the job state, notifying the user and preparing the download URLs.

A synchronized copy of the current job queue is permanently stored on the system's file storage. In the case of a system failure, the queue is automatically restored from that file. It is also possible to manually stop the queue without terminating it, e.g. for server administration. In this case, the currently running job will be regularly processed, but further jobs are delayed until the queue is put into the running state again. It is also possible to schedule new jobs.

##### 1.2.4 Job management and results download

After submission, the job is queued in the job queue and can be managed in the "Jobs" section. The job management page is divided into four parts: queued jobs, running jobs, finished jobs and the job history. The jobs are listed in chronological order along with a waiting status indicating the position in the job queue. A single user can queue up to three jobs at a time (i.e. no more than three jobs can be queued waiting to be processed by a single user, however, there is no limit to the total number of jobs for a user).

**Supplementary Figure 2** shows a screenshot from an exemplary job progress. Once a job is actively being processed, the user can monitor the current progress and view information about the job, such as the elapsed runtime, warning and error messages. When a job is regularly completed, it is moved to the finished jobs section from where the result files can be downloaded. A job can also be canceled, in which case it is stopped and also moved to the finished jobs section.

For each successfully finished job the web service generates a `.scores` file that contains the $n$

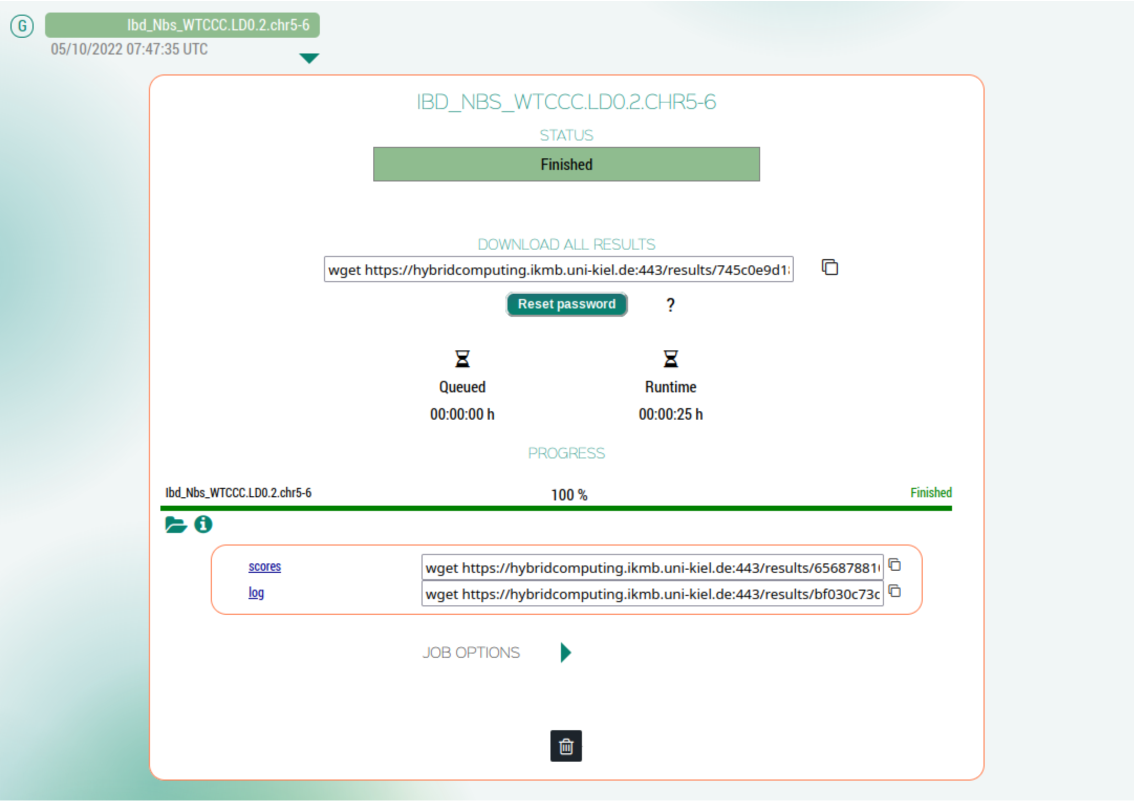

Supplementary Figure 3: Screenshot of a finished job with exemplary result files in *GWAIS-Web*.

best results of the analysis according to the first selected test method, whereby $n$ was also selected by the user. The score file is provided as a zipped `.gz` file. In addition, the *HybridGWAIS* `.log` log file is provided. **Supplementary Figure 3** shows a screenshot from a finished job with exemplary result files.

The result files are registered in our database with a unique random string for each file. The web service then generates a secure download URL for each file over an encrypted *https*-connection based on this random string. We use the *rewrite engine* by the Apache server to decode this URL to redirect a download request to our download engine, and the download engine queries the database for the associated file.

The user may download each result file separately from the web browser, but an individual script to download all available files at once from a (Linux) command line terminal is also provided to the user. For this purpose, the user receives a notification email for all completed, cancelled or otherwise terminated jobs, containing a generated password uniquely associated with that job. The files are locked by default and a direct download via the file URL is only possible when the user is either logged-in to download individual files directly from the website or the received password is entered via the download script on the command line to authenticate the download. Technically, the password has to be sent as a *POST* command after establishing the encrypted *https* connection to the web server to download a file, which is done automatically by the provided script. This is advantageous if the result files have to be analyzed or post-processed on a different system than the user's computer.

Note that we automatically delete all data related to a job (with the exception of status and log files) from our server 7 days after the job has finished unless the job is manually deleted by the user before. We keep status and log files together with the job metadata for the user in the job history section. Of course, jobs in the job history can also be deleted manually (either individual jobs or all deleted jobs at once). Anyway, jobs older than one year are automatically removed completely from the history.

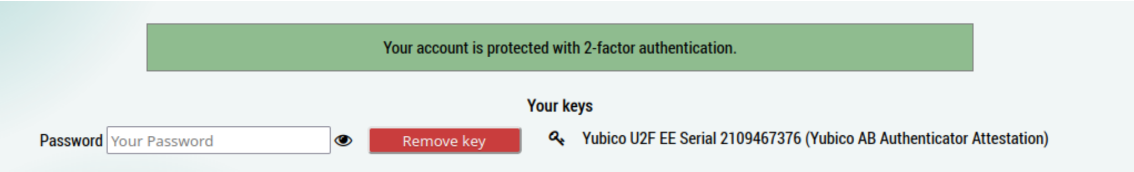

Supplementary Figure 4: Screenshot of exemplary account settings with a registered key device for 2-factor authentication.

##### 1.2.5 Account management

As stated above, a valid email address and a password are required to set up a personal user account. The password is not stored in plain text, but as a secure SHA-256 hash on our server. Users are able to change their email address and password at any time in the account management section.

*GWAIS-Web* also provides a password recovery function to set a new password in the case the user has forgotten the login password. The user can click on the link "Forgot Password?" next to the password field and enter the email address and a captcha code displayed as image, which is necessary to prevent abuse by bots. If the captcha code is correct and the email address is registered in our database, a link to reset the password is sent to the user. After clicking the link, a new random password is sent to that email address. The user can now sign in again and change the password.

Optionally, to improve account security, the user can activate 2-factor authentication for her/his account based on the recent standard *Web Authentication API (Webauthn)* [5]. Webauthn enables usage of hardware authenticators with public and private key-based credentials to perform an SSL handshake between server and the client's authenticator as trusted device. A trusted device may be a USB key dongle, fingerprint reader or facial recognition on a phone or any other applicable device. In detail, an account which is protected by 2-factor authentication requires the correct password and the correct authentication of one of the registered trusted devices to verify the user's identity upon login. Note, that after the registration of a trusted device, the login is not possible from clients where the device is not available, e.g. if a smartphone's integrated fingerprint reader is used as trusted device, the login is not possible from devices other than that smartphone. To avoid this problem, the user is able to register more than one trusted device. We recommend to use at least one portable device, such as a USB key dongle, for 2-factor authentication. Note that once 2-factor authentication has been activated, it is no longer possible to log in without a trusted device. **Supplementary Figure 4** shows an exemplary registered trusted device in the account settings.

The user may also delete her/his account in the account settings, which results in an immediate deletion of all data associated with this account from our server without exception. Specifically, the login credentials are removed from our database together with all uploaded files and data created for this user as a result from using the service.

### 2 Basic concepts of the *HybridGWAIS* software

#### 2.1 Contingency tables

For any variant pair $(A, B)$ a contingency table represents the number of samples in a dataset that carry a specific genotype information. In particular, an entry $n_{ij}$ represents the number of samples that carry the information $g_A = i$ at variant $A$ and $g_B = j$ at variant $B$. Thus, a contingency table for pairwise genotypic tests contains $3 \times 3$ entries. Since we are focusing on binary traits, we require a contingency table for each state, w.l.o.g. one for the *case* and one for the *control* group, respectively, and denote their entries by $n_{ij}^{\text{case}}$ and $n_{ij}^{\text{ctrl}}$ (see **Supplementary Figure 5**).

| cases ($Y=1$) | | variant $A$ | | |
|---|---|---|---|---|
| | | 0 | 1 | 2 |
| variant $B$ | 0 | $n_{00}^{\text{case}}$ | $n_{01}^{\text{case}}$ | $n_{02}^{\text{case}}$ |
| | 1 | $n_{10}^{\text{case}}$ | $n_{11}^{\text{case}}$ | $n_{12}^{\text{case}}$ |
| | 2 | $n_{20}^{\text{case}}$ | $n_{21}^{\text{case}}$ | $n_{22}^{\text{case}}$ |

| controls ($Y=0$) | | variant $A$ | | |
|---|---|---|---|---|
| | | 0 | 1 | 2 |
| variant $B$ | 0 | $n_{00}^{\text{ctrl}}$ | $n_{01}^{\text{ctrl}}$ | $n_{02}^{\text{ctrl}}$ |
| | 1 | $n_{10}^{\text{ctrl}}$ | $n_{11}^{\text{ctrl}}$ | $n_{12}^{\text{ctrl}}$ |
| | 2 | $n_{20}^{\text{ctrl}}$ | $n_{21}^{\text{ctrl}}$ | $n_{22}^{\text{ctrl}}$ |

Supplementary Figure 5: Contingency tables for cases and controls. $n_{ij}$ reflect the number of occurrences for the corresponding genotype combination in a given pair of variants.

#### 2.2 Test methods

Currently, *HybridGWAIS* supports the test methods listed in **Supplementary Table 1**. In addition, the user may add the calculation of linkage disequilibrium ($r^2$) to the selected methods. It is also possible to enable filtering based on LD on-the-fly. All methods are described in the following.

Supplementary Table 1: Currently supported test methods available in *HybridGWAIS* and *GWAIS-Web*.

| **2nd-order (pairwise) methods** | **3rd-order methods** |
|---|---|
| logistic regression | logistic regression |
| BOOST | |
| log-linear test | |
| mutual information | mutual information |
| information gain | information gain |
| **optionally appendable to at least one selected method:** | |
| linkage disequilibrium ($r^2$) | linkage disequilibrium (all pairwise $r^2$) |

##### 2.2.1 Logistic regression

The logistic regression test implemented in *HybridGWAIS* is taken from PLINK's [6] multiplicative logistic regression model, with $\beta_3$ indicating the interaction effect:

$$\ln \frac{P(Y=1|X_A=g_A, X_B=g_B)}{P(Y=0|X_A=g_A, X_B=g_B)} = \beta_0 + \beta_1 g_A + \beta_2 g_B + \beta_3 g_A g_B \tag{1}$$

$Y$ defines the categorization if a sample is *case* ($Y=1$) or *control* ($Y=0$). $X_A$ and $X_B$ represent random variables correlated with the observation of genotypes at variants $A$ and $B$, respectively. The possible outcomes of $X_{A/B}$ are $g_{A/B} \in \{0,1,2\}$ representing the observed genotype (0 = homozygous reference, 1 = heterozygous, 2 = homozygous variant).

The model is now fitted using the contingency tables generated in Sect. 2.1 and an iterative Newton method. For the first iteration, we start with $\boldsymbol{\beta} = (0,0,0,0)$. The iterations are then processed as follows:

1. For each sample $i$, compute intermediate variables

$$p_{ij}^{(t)} = \left(1 + e^{-\left(\beta_0^{(t)} + i\beta_1^{(t)} + j\beta_2^{(t)} + ij\beta_3^{(t)}\right)}\right)^{-1} \tag{2}$$

$$p_{ij}^{(t),\text{ctrl}} = p_{ij}^{(t)}, \quad p_{ij}^{(t),\text{case}} = p_{ij}^{(t)} - 1, \quad v_{ij}^{(t)} = p_{ij}^{(t)}\left(1 - p_{ij}^{(t)}\right)\left(n_{ij}^{\text{case}} + n_{ij}^{\text{ctrl}}\right) \tag{3}$$

2. Compute gradient

$$\boldsymbol{\nabla}^{(t)} = \left(\sum_{ij} N_{ij}^{(t)}, \sum_{ij} iN_{ij}^{(t)}, \sum_{ij} jN_{ij}^{(t)}, \sum_{ij} ijN_{ij}^{(t)}\right) \tag{4}$$

where

$$N_{ij}^{(t)} = \left(n_{ij}^{\text{case}} p_{ij}^{(t),\text{case}} + n_{ij}^{\text{ctrl}} p_{ij}^{(t),\text{ctrl}}\right) \tag{5}$$

3. Compute symmetric Hessian matrix

$$\boldsymbol{H}^{(t)} = \left(h_{pq}^{(t)}\right)_{p,q=0}^{3} = \begin{pmatrix} \sum v_{ij}^{(t)} & & \cdots & \\ \sum i v_{ij}^{(t)} & \sum i^2 v_{ij}^{(t)} & & \vdots \\ \sum j v_{ij}^{(t)} & \sum ij v_{ij}^{(t)} & \sum j^2 v_{ij}^{(t)} & \\ \sum ij v_{ij}^{(t)} & \sum i^2 j v_{ij}^{(t)} & \sum i j^2 v_{ij}^{(t)} & \sum i^2 j^2 v_{ij}^{(t)} \end{pmatrix} \tag{6}$$

where each sum is evaluated over all indexes $i$ and $j$

4. Compute $\boldsymbol{\Delta\beta}^{(t)} = \left(\Delta\beta_j^{(t)}\right)_{j=0}^{3}$ by efficiently solving the linear system

$$\boldsymbol{L}^{(t)} \boldsymbol{L}^{(t)T} \boldsymbol{\Delta\beta}^{(t)} = \boldsymbol{\nabla}^{(t)} \tag{7}$$

using the Cholesky decomposition $\boldsymbol{L}^{(t)} = \left(l_{jk}^{(t)}\right)_{j,k=0}^{3}$ of $\boldsymbol{H}^{(t)}$ with

$$l_{jk}^{(t)} = \begin{cases} 0 & \text{if } k > j \\ \sqrt{h_{jj}^{(t)} - \sum\limits_{s=1}^{j-1} l_{js}^2} & \text{if } k = j \\ \frac{1}{l_{kk}} \left(h_{jk}^{(t)} - \sum\limits_{s=1}^{k-1} l_{js} l_{ks}\right) & \text{if } k < j \end{cases} \tag{8}$$

5. Update model parameters

$$\boldsymbol{\beta}^{(t+1)} \leftarrow \boldsymbol{\beta}^{(t)} - \boldsymbol{\Delta}\beta^{(t)} \tag{9}$$

If $\sum_j \Delta\beta_j^{(t)}$ approaches zero, i.e. there is no more significant change ($\leq 0.0001$), the process stops with $\boldsymbol{\beta}^{(t+1)}$ as the current result. Otherwise, the next iteration is started with step 1. However, if the change does not converge to zero, the process stops after a fixed number of iterations (currently 16 which was taken over from PLINK).

The result of the logistic regression test in PLINK is composed of three components, namely the test statistic, its approximate p-value and the odds-ratio. The test statistic $\chi^2$ is calculated as

$$\chi^2 = \frac{\beta_3^2}{\varepsilon^2}. \tag{10}$$

$\varepsilon$ is the standard error for the $g_A g_B$-term in Eq. 1. It can directly be determined by solving the linear system $\boldsymbol{H}^{(t)}\boldsymbol{e} = (0,0,0,1)$ and defining $\varepsilon^2 = e_3$.

Accordingly, it follows

$$\varepsilon^2 = \frac{1}{\left(l_{33}^{(t)}\right)^2}. \tag{11}$$

The test statistic is assumed to follow a chi-squared distribution $\chi_1^2$ with one degree of freedom. Accordingly, the p-value can directly be approximated from the respective cumulative distribution function (CDF):

$$\text{Pval}(x) = 1 - \text{CDF}(\chi_1^2(x)) \tag{12}$$

Finally, the odds-ratio is defined as

$$\text{OR} = e^{\beta_3}. \tag{13}$$

For a third-order logistic regression test the following multiplicative logistic regression model is fitted analogue to the procedure described above:

$$\ln \frac{P(Y=1|X_A=g_A,X_B=g_B,X_C=g_C)}{P(Y=0|X_A=g_A,X_B=g_B,X_C=g_C)} = \beta_0+\beta_1 g_A+\beta_2 g_B+\beta_3 g_C + \beta_4 g_A g_B + \beta_5 g_A g_C + \beta_6 g_B g_C + \beta_7 g_A g_B g_C \quad (14)$$

The same definitions as above apply plus $X_C$ and $g_C$ describe the random variable and its outcomes of the third variant $C$. The reported interaction effect is taken from $\beta_7$.

##### 2.2.2 BOOST and log-linear test

In BOOST [7] an interaction is defined as the difference between the log-likelihoods of the saturated model $\hat{L}_S$ and the homogenous model $\hat{L}_H$. As no closed solution for the homogenous model exists, the BOOST method was divided into two parts.

**BOOST pre-filter:** At first, the *Kirkwood Superposition Approximation (KSA)* of the homogenous model $\hat{L}_{KSA}$ is calculated. Wan et al. showed that the difference of the log-likelihoods is an upper bound to the desired calculation of the interaction effect:

$$\hat{L}_S - \hat{L}_H \leq \hat{L}_S - \hat{L}_{KSA} \quad (15)$$

This correlation is used as a pre-filter, so the log-linear test (i.e. fitting of the homogeneous model) is done only when the pre-filter exceeds a certain threshold $\tau$, which is fixed to $\tau = 15$ according to [7]:

$$\hat{L}_S - \hat{L}_{KSA} \overset{!}{\geq} \tau \quad (16)$$

The approximated interaction effect is then calculated by:

$$\hat{L}_S - \hat{L}_{KSA} = n \sum_{ijY} \left[ \hat{\pi}_{ijY} \log \frac{\hat{\pi}_{ijY}}{\hat{p}^K_{ijY}} \right] \quad (17)$$

where $\hat{\pi}_{ijY}$ is the joint distribution of the saturated model, which can be replaced by the observed relative probability of a certain genotype combination $ij$ for a certain trait $Y$ ($N$ is the number of samples):

$$\hat{\pi}_{ijY} = \frac{n^Y_{ij}}{N} \quad (18)$$

Further, $\hat{p}^K_{ijY}$ is the KSA of the distribution obtained under the homogeneous association model and can be computed as:

$$\hat{p}^K_{ijk} = \frac{1}{\eta} \frac{\pi_{ij\cdot}\pi_{i\cdot k}\pi_{\cdot jk}}{\pi_{i\cdot\cdot}\pi_{\cdot j\cdot}\pi_{\cdot\cdot k}} \quad (19)$$

whereby

$$\eta = \sum_{ijY} \frac{\pi_{ij\cdot}\pi_{i\cdot Y}\pi_{\cdot jY}}{\pi_{i\cdot\cdot}\pi_{\cdot j\cdot}\pi_{\cdot\cdot Y}}. \quad (20)$$

(Note, we use the dot notation to indicate a sum over a subscript, e.g. $\pi_{ij\cdot} = \sum_Y \pi_{ijY}$ and $\pi_{\cdot\cdot Y} = \sum_{ij} \pi_{ijY}$.)

**Log-linear test:** If the approximated interaction effect using the KSA pre-filter exceeds the predefined threshold of $\tau = 15$, BOOST computes the log-linear test to obtain a better approximation to the desired interaction effect using the homogeneous model $\hat{L}_S - \hat{L}_H$. In *HybridGWAIS*, the user may choose the log-linear test as a separate method to directly calculate the interaction effect without the BOOST pre-filter based on the KSA.

The log-linear test is computed using *Iterative Proportional Fitting* of the homogenous association model using a maximum of 34 iterations until the error is below $\varepsilon^{(t)} \leq 0.001$. The process starts at iteration $t = 1$ and is initialized with $\mu^{(0)}_{ijY} = 1$.

1. Compute intermediate $\mu'_{ijY}$ for all $ijY$:

$$\mu'^{(t)}_{ijY} = \mu^{(t-1)}_{ijY} \frac{n^{\cdot}_{ij}}{\mu^{(t-1)}_{ij\cdot}} \tag{21}$$

2. Compute intermediate $\mu''_{ijY}$ for all $ijY$:

$$\mu''^{(t)}_{ijY} = \mu'^{(t)}_{ijY} \frac{n^{Y}_{i\cdot}}{\mu'^{(t)}_{i\cdot Y}} \tag{22}$$

3. Compute $\mu_{ijY}$ for all $ijY$:

$$\mu^{(t)}_{ijY} = \mu''^{(t)}_{ijY} \frac{n^{Y}_{\cdot j}}{\mu''^{(t)}_{\cdot jY}} \tag{23}$$

4. Compute error $\varepsilon$:

$$\varepsilon^{(t)} = \sum_{ijY} \left| \mu^{(t)}_{ijY} - \mu^{(t-1)}_{ijY} \right| \tag{24}$$

The interaction score is finally calculated as:

$$\hat{L}_S - \hat{L}_H \approx 2 \sum_{ijY} n^Y_{ij} \log \frac{n^Y_{ij}}{\mu^{(t)}_{ijY}} \tag{25}$$

*HybridGWAIS* reports the interaction score with the last error $\varepsilon^{(t)}$ and an approximated p-value for the interaction score under the assumption of a $\chi^2$-distribution with four degrees of freedom.

Note that BOOST and the log-linear test are not yet implemented as third-order methods in *HybridGWAIS*.

##### 2.2.3 Entropy-based tests

Entropy-based tests in *HybridGWAIS* include *mutual information* and *information gain* (which is also known as *interaction information*) [8, 9]. By defining the observations of the genotypes in a marker pair or triple and the corresponding phenotypes as random variables, one can easily calculate their *entropy* $H$, which forms the basis of the methods described below. For a random variable $X$ and its possible outcomes $\{x_i\}$ the entropy is defined as

$$H(X) = -\sum_i p(x_i) \log p(x_i). \tag{26}$$

**Mutual information:** The first method computes the *mutual information (MI)* between the observed genotypes and the corresponding phenotype. Let $X_1$ be the random variable for observations of genotypes at the first marker in a pair, and $X_2$ for the second marker. For third-order tests, let $X_3$ be the random variable for the third marker. In general, let $Y$ be the random variable for the phenotype, i.e. either case or control. Mutual information describes the overlap of the combined entropies for the genotypes and the entropy for the phenotype, i.e. for pairs

$$I(X_1, X_2; Y) = H(X_1, X_2) + H(Y) - H(X_1, X_2, Y) \tag{27}$$

and for triples

$$I(X_1, X_2, X_3; Y) = H(X_1, X_2, X_3) + H(Y) - H(X_1, X_2, X_3, Y) \tag{28}$$

respectively.

**Information gain:** The second method computes the *information gain (IG)* from the observed genotypes to the corresponding phenotype. It is based on the mutual information of the single random variables to the phenotype and the combined mutual information as calculated above in Eqs. 27 and 28. *HybridGWAIS* calculates IG according to the definition of Jakulin et al. [8, 9], i.e. for pairs

$$I(X_1; X_2; Y) = I(X_1, X_2; Y) - I(X_1; Y) - I(X_2; Y) \tag{29}$$

and for triples

$$\begin{aligned} I(X_1; X_2; X_3; Y) = I(X_1, X_2, X_3; Y) - I(X_1, X_2; Y) - I(X_1, X_3; Y) - I(X_2, X_3; Y) \\ + I(X_1; Y) + I(X_2; Y) + I(X_3; Y) \end{aligned} \tag{30}$$

respectively. Note that information gain may be negative.

##### 2.2.4 Linkage disequilibrium

Linkage disequilibrium is usually measured as an $r^2$-score and is a measure of similarity between two variants. It is defined as

$$r^2 = \frac{D^2}{p_A(1-p_A)p_B(1-p_B)} \quad \text{with} \quad D = p_{AB} - p_A p_B. \tag{31}$$

$D$ is the distance between the observed allele frequency $p_{AB}$ at loci $A$ and $B$ and the expected allele frequency $p_A p_B$ assuming statistical independence. Thus, $r^2$ is a normalized measure for $D$ which can be used for comparison of different variant pairs. The allele frequencies $p_A$ and $p_B$ can directly be determined as

$$p_A = \frac{2n_{00} + 2n_{10} + 2n_{20} + n_{01} + n_{11} + n_{21}}{2N} \tag{32}$$

and

$$p_B = \frac{2n_{00} + 2n_{01} + 2n_{02} + n_{10} + n_{11} + n_{12}}{2N}, \tag{33}$$

respectively, where $n_{ij} = n_{ij}^{\text{case}} + n_{ij}^{\text{ctrl}}$ for all $i$, $j$. Unfortunately, the determination of the allele frequency $p_{AB}$ from genotypic data is not trivial. This is due to the unknown phase when two heterozygous genotypes face each other in a variant pair. Basically, it can be defined as

$$p_{AB} = \frac{2n_{00} + n_{01} + n_{10} + x}{2N} \tag{34}$$

with $x$ meeting $x \leq n_{11}$. $x$ has to satisfy the following equation whose solution is omitted here for simplicity:

$$(f_{00} + x)(f_{11} + x)(n_{11} - x) = (f_{01} + n_{11} - x)(f_{10} + n_{11} - x)x \tag{35}$$

where $f_{ij}$ is the number of allele combinations $ij$ we know for sure, e.g. $f_{00} = 2n_{00} + n_{01} + n_{10}$ and $f_{11} = 2n_{22} + n_{21} + n_{12}$.

Note that in general, there exist more than one solution for this equation. Thus, *HybridGWAIS* calculates and reports the smallest and the largest of the (at maximum three) possible solutions for $r^2$. If an LD-filter is applied in addition to the test calculation, only pairs with the largest solution for $r^2$ being below the filter threshold are reported.

For third-order tests the $r^2$-score is computed for all three pairwise combinations in a variant triple. In contrast to the pairwise method only the largest solution of each of the pairwise calculations is reported. If an LD-filter is applied to third-order methods, triples only pass the filter if all pairwise $r^2$-scores are below the filter threshold.

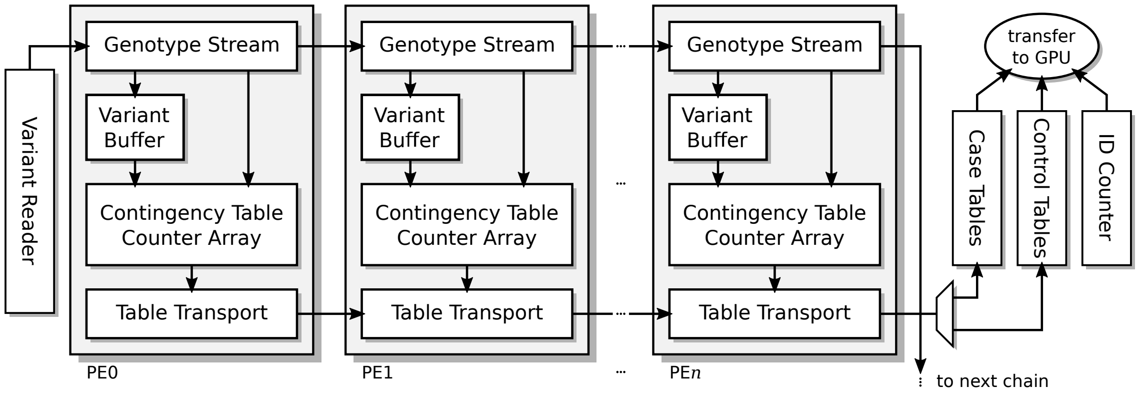

Supplementary Figure 6: Chain of processing elements (PEs) in the FPGA design for creating contingency tables. Our design for the *Xilinx Kintex KU115* FPGA contains 480 PEs which allows the creation of 480 contingency tables in parallel.

### 3 FPGA and GPU acceleration in *HybridGWAIS*

#### 3.1 FPGA-based creation of contingency tables

The acceleration method of combining FPGAs and GPUs is divided into two main parts. Firstly, the FPGA accelerator creates the contingency tables for each marker pair or triple, while, secondly, the GPU accelerator processes each contingency table by applying the selected statistical tests. For a complete exhaustive analysis of all marker pairs (or triples) we refer to the method description in [10] and [11]. We shortly summarize the method for the creation of pairwise contingency tables here. The modifications required to handle arbitrary region selections are described below in Section 3.4.

The FPGA design targets the *Xilinx Kinex UltraScale KU115* FPGAs with 16 GB of attached DRAM (distributed over two DRAM modules) on our *Alpha Data ADM-PCIE-8K5* accelerator cards. The design sustains a pipeline consisting of a chain of 480 process elements (PEs) divided into two subchains with 240 PEs each. After a short initialization phase, the chain produces 480 contingency tables concurrently while the genotype data of one marker is streamed through the pipeline at a speed of 266 MHz and 8 genotypes per clock cycle. This sums up to a peak performance of about 20.4 million contingency table pairs per second for an exemplary dataset containing about 50,000 samples.

In detail, each PE in a chain is organized into a memory buffer to store the genotypes of all samples at a single marker position, implemented in local block RAM (BRAM), and a counter array for the contingency table entries implemented in registers (see **Supplementary Figure 6**). The PE is able to receive a stream of genotype data from a previous PE and to process it while also delivering it to the next PE. For each PE the local genotype buffer is filled with the first incoming genotype data for one complete marker. In particular, the data for the first marker is only stored in the local memory and not forwarded to the next PE. Subsequent variants are then forwarded and simultaneously used to form genotype pairs with the data stored in the local memory. Dependent on the current pair, the corresponding counter of the contingency table is incremented. By streaming eight genotypes at once for each marker pair, i.e. incrementing the corresponding counters of eight genotype combinations at the same time, we could achieve a high throughput of genotype data in each PE.

The genotype data for the analysis is stored in the first half of the attached DRAM. Streaming is organized by sending the genotypes of the case samples at the current variant position first, followed by the genotypes of the control samples. Thus, the contingency tables for cases and controls are alternately generated using the same logic resources in the PE. Currently, our design supports a maximum of 65,535 cases or controls, implying 16 bit counters for the entries of the contingency table and block RAM (BRAM) resources for the memory buffer of size 128 kbit.

The contingency tables are retrieved from the PEs again via the PE chain. After all genotypes from either the cases or the controls were streamed, the table is provided to the next PE in one clock cycle. Each PE hands over incoming tables to the next PE first before it sends its own table. All tables are collected at the end of the chain in a separate buffer (implemented in the second half of the FPGA-attached DRAM) before they are transmitted to the GPU via the host.

Thus, the collection of tables requires as many clock cycles as there are PEs in the chain implying that the minimum number of either cases or controls for the most efficient utilization of the chain is exactly eight times the number of PEs in the chain. In order to prevent delays resulting from collecting the contingency tables, we divided the complete chain into subchains with continuous genotype streaming but separate table collection units.

#### 3.2 GPU-based creation of contingency tables

In the case the FPGA acceleration is disabled or not available the contingency tables can be created directly on the GPU. The host loads the genotype database in the GPU local memory beforehand and provides a continuous flow of buffers to the GPU that contain index pairs indicating which marker pairings are to be analyzed next. For each buffer a number of CUDA threads is launched, each thread processing one pair.

The CUDA thread firstly creates the contingency table by reading the required genotype data from local GPU memory and then continues the calculation of the test statistic in the same way as with an attached FPGA accelerator (see next Section 3.3). There is no need to store the contingency table anywhere other than in the local thread registers. Thus, we only create warp divergence while reading the individual marker data for each thread, but due to the packed variant-wise data format with a 2-bit encoding per genotype, each thread only requires a few kilobytes which can be processed almost immediately. Furthermore, in most cases, the marker pairs in the current warp share the first marker and the second markers in the pairs are in consecutive order. Together with the word-aligned genotype data, this is an optimal scenario for fast access to the local GPU memory.

#### 3.3 GPU-based computation of statistical tests

In the hybrid solution with combined FPGA and GPU accelerators, the buffers from the FPGA containing contingency tables are transferred to local GPU memory. We use a default transmission buffer size of 256 MB which may hold up to 6.7 million table pairs. In a GPU-only acceleration scenario, the transmission buffers contain only index pairs for which the GPU needs to create the contingency tables from genotype data in the local GPU memory itself (as described above in Section 3.2).

The computation process follows a simple parallelization scheme over CUDA threads. By setting the block size to the maximum supported block size and the grid size to evenly distribute the contingency tables over the blocks, each thread processes exactly one contingency table pair, and only one kernel call per buffer is required. Besides the distribution of the contingency table data and the writeback of the results no calls to local GPU memory are required from a GPU thread in the hybrid scenario (in the GPU-only scenario each thread needs to recall the required genotype data from local memory).

The test methods are implemented as described in Section 2.2 using double precision floating point formats where applicable. (In contrast, PLINK uses only the single precision floating point format in its computations.) The resulting test scores are written into a result buffer. We provide one result buffer for each table transmission buffer, which is transferred to the host as soon as processing the table buffer has finished.

#### 3.4 Chromosomal region selection

##### 3.4.1 Region selection options

In *HybridGWAIS* the user is able to select arbitrary regions to reduce the analysis space by choosing intervals in the input dataset for each variant marker separately. The space usually opens up to $\binom{n}{2} = n(n-1)/2$ pairwise combinations ($n$ is the number of variants) for exhaustive pairwise tests, and $\binom{n}{3} = n(n-1)(n-2)/6$ triple combinations for exhaustive third-order tests. For pairwise tests, users are allowed to concentrate on certain regions by selecting an interval subset for the first marker and another interval subset for the second marker. A third interval may be specified for the third marker when doing a third-order test.

Additionally, users may define an additional interval for a region that they wish to exclude from the analysis. This *exclude region* may be applied either to the complete dataset or only to the interval selection for the first marker. The latter option may be useful if a user wants to test the whole dataset excluding a certain region against the whole data but including that region.

According to the user selections the software simply determines overlapping and non-overlapping regions from the selected intervals. It then determines which tests have to be evaluated and which not. The implementation differs for the selected type of computation whether or not FPGA-acceleration is used (see the following Sects. 3.4.2 and 3.4.3).

##### 3.4.2 Processing of regions with CPU-only or GPU-only acceleration

If FPGA-acceleration is disabled and the computation is evaluated only on the CPU cores or using GPU-acceleration, the loops that circulate through the number of all tests are programmed to simply leave out regions that are not selected by the user. For CPU-only processing this means that only function calls to variant pairs/triples of interest are performed leaving out unselected combinations. For GPU-acceleration, the host creates lists of pairs/triples to be tested on the GPU anyway, which are then provided to the GPU via the transmission buffers. With enabled region selections the host simply adds only pairs/triples of interest to this list.

##### 3.4.3 Processing of regions with FPGA+GPU acceleration

As the FPGA pipeline is designed to concurrently generate contingency tables from a continuous flow of variant data, region selections have to be handled differently when compared to CPU-only or GPU-accelerated processing. The original pipeline designed in [10] was not able to handle region selections. After transmitting the complete dataset to the FPGA-attached local DRAM the pipeline controller starts streaming the complete data through the pipeline. In the first cycle, this generates the contingency tables for the first 480 variants (corresponding to the number of PEs in the pipeline) against all variants in the dataset. In the second cycle, the controller streams the data again, but leaves out the first 480 variants. This effectively generates the contingency tables for the next 480 variants against the remainder. This process continues until the controller cannot start another cycle due to having to leave out all available variants.

In order to handle region selections we modified the pipeline controller in two ways. First, the pipeline cycles can be stopped if the variant to begin a new cycle exceeds a defined index. Second, the controller can "jump" to a certain variant index after the initialization phase of a pipeline cycle. With this modifications the pipeline is able to handle not only regions that completely overlap, but also regions that partially overlap in the beginning or do not overlap at all. As the host system determines these areas of overlap and non-overlap in advance, the FPGA-accelerator can now handle any arbitrary region selections in several runs. Due to the pipeline nature these modifications imply that some contingency tables may be created that are not required for the selected analysis, especially in the initialization phase and in the last pipeline cycle. The tables are still handed over to the GPU-accelerator that computes the selected test statistics, but the results are filtered afterwards by the host. This overhead in computation may have an impact in the runtime, especially if inefficient regions are chosen (e.g. selecting only a single variant in a dataset to be tested against the rest), but in the general case, the impact will be negligible and in most cases the analysis speed will still be faster in comparison to a GPU-only acceleration. Anyway, when using *HybridGWAIS* in our web service *GWAIS-Web*, the runtime prediction computes both expected runtimes (with and without using FPGA-acceleration) and eventually starts the analysis with the smaller predicted runtime.

##### 3.4.4 Proximity exclude range

In addition to the chromosomal region selection abilities presented above, the user may also define a *proximity exclude range*. This range is measured in base pairs and defines an area around every marker where tests should be omitted if the second (or third) marker resides within that range. For example, a proximity exclude range of 50 kbp omits all tests where the two markers in a pair (or any two markers in a triple) are located in proximity closer than 50,000 base pairs. Similar to the region selection options, the implementation differs for the selected acceleration type. For CPU-only or GPU-accelerated processing the distance is checked before running the test, which effectively skips unwanted tests beforehand. In contrast for FPGA-accelerated runs, the tests are evaluated first and the distance of the pairs is checked afterwards excluding unwanted test results before adding them to the result list.

#### 3.5 Data collection and post processing

Multiple threads on the host system perform the collection and post-processing of results. Results are sorted by the result score of the first user selected test method. For this purpose, a *min-max heap* data structure with a user definable size limit is used. Alternatively, the user may define a significance threshold which prevents results not exceeding this limit not to be inserted into the result heap in the first place.

Each thread keeps its own instance of a min-max heap to avoid lock conditions. After processing all transmission buffers, the results are merged into a single instance. The final results are written into a tab-delimited table file containing the marker IDs, indices and all available scores.

#### 3.6 Design limitations

In general, *HybridGWAIS* may process an unlimited amount of genotype data as long as it fits into the host memory. Additional memory is only required for the result heaps whose size depends on the number of desired results selected by the user. However, if GPU acceleration is requested, the genotype data must fit into the local GPU memory together with some result transfer buffers, which still does not state a real problem with up-to-date graphics accelerators compared to the required runtime if an analysis should be finished in feasible time.

The FPGA accelerator though has some technical limits which might affect the analysis in rare cases. Currently, our FPGA-design supports a maximum of 131,070 samples (whereby neither cases nor controls may exceed a maximum number of 65,535) due to the trade-off we had to face in distributing the available local block RAM to gain a maximum number of processing elements. We managed to implement 480 PEs in total divided into two subchains with 240 PEs each, as already mentioned in Section 3.1. Thus, if a dataset contains more than the allowed number of samples, *HybridGWAIS* automatically switches to GPU-only acceleration in order to still be able to run the analysis. We are currently working on a solution that allows to distribute the data into several runs over the number of samples.

Furthermore, for a most efficient use of the FPGA-accelerator, the input data should contain at least 1920 cases or controls. This limitation results from the fact that the PE-chain requires 240 clock cycles (corresponding to the number of PEs in a subchain) to fetch the contingency tables from each PE, and that in each clock cycle 8 genotypes are processed concurrently. However, if a dataset contains less cases or controls, *HybridGWAIS* fills up the data with a padding consisting of unknown genotypes.

Likewise, user selected chromosomal regions of less than 480 variants in a single interval will most likely generate a slight computational overhead in *HybridGWAIS* (see Sect. 3.4.3). For this reason, our web service *GWAIS-Web* compares the runtime prediction of *HybridGWAIS* with FPGA+GPU acceleration to GPU-only acceleration and chooses the configuration with the smaller predicted runtime.

As the genotype data for the analysis is stored in one half of the FPGA-attached DRAM, an FPGA-accelerated run is also limited by a maximum of 8 GB of genotype data. However, we overcome this limitation easily by dividing up the analysis space equally into several runs (using the priniciples described in Section 3.4 above), where the data required for each run fits into the limitations.

### 4 Supplementary benchmark information

#### 4.1 Creation of benchmark datasets

For the benchmark data we firstly generated a large dataset of 1 million simulated individual samples based on the approximately 3 million variants of chromosome 1 from the *Haplotype Reference Consortium (HRC)* reference panel *HRC r1.1* freely available at `ftp://ngs.sanger.ac.uk/production/hrc/HRC.r1-1/HRC.r1-1.GRCh37.wgs.mac5.sites.vcf.gz`. For each sample we randomly generated the alleles for each variant according to the allele frequency presented in this file. We converted the resulting VCF file to PLINK's bed/bim/fam format and randomly sampled subsets with different sizes (a different number of variants and samples) from this file using PLINK 1.9. The resulting files were then used for our benchmarks.

#### 4.2 Supplementary benchmark results

Supplementary Table 2: Wall-clock runtimes in seconds of PLINK's epistasis test with a different number of variants ($M$) and a different number of samples ($N$) using 32 computing threads on our benchmark system.

| | $N$ = 1000 | 5000 | 10,000 | 50,000 | 100,000 | 500,000 | 1,000,000 |
|---|---|---|---|---|---|---|---|
| $M$ = 10,000 | 16.05 | 125.41 | 290.34 | 3315.08 | 13898.12 | 192001.73 | 443975.95 |
| 30,000 | 146.34 | 1129.92 | 2605.55 | 30428.37 | 127807.41 | - | - |
| 50,000 | 473.26 | 3719.40 | 8776.58 | 83003.27 | - | - | - |

Supplementary Table 3: Wall-clock runtimes in seconds of *HybridGWAIS* logistic regression test with a different number of variants ($M$) and a different number of samples ($N$) using CPU-only computation with 32 computing threads on our benchmark system.

| | $N$ = 1000 | 5000 | 10,000 | 50,000 | 100,000 | 500,000 | 1,000,000 |
|---|---|---|---|---|---|---|---|
| $M$ = 10,000 | 21.73 | 64.14 | 119.57 | 558.84 | 1106.14 | 5488.78 | 10956.37 |
| 30,000 | 146.32 | 545.35 | 1041.82 | 4970.56 | 9872.64 | 49213.27 | - |
| 50,000 | 400.78 | 1504.99 | 2869.07 | 13802.26 | 27410.12 | - | - |
| 100,000 | - | - | 11527.64 | 55044.75 | - | - | - |
| 300,000 | - | - | 103520.70 | - | - | - | - |

Supplementary Table 4: Wall-clock runtimes in seconds of *HybridGWAIS* logistic regression test with a different number of variants ($M$) and a different number of samples ($N$) using GPU acceleration with an Nvidia Tesla P100 GPU on our benchmark system.

| | $N$ = 1000 | 5000 | 10,000 | 50,000 | 100,000 | 500,000 | 1,000,000 |
|---|---|---|---|---|---|---|---|
| $M$ = 10,000 | 11.80 | 9.22 | 9.64 | 22.54 | 39.07 | 171.79 | 333.26 |
| 30,000 | 18.69 | 24.67 | 36.34 | 145.06 | 283.74 | 1397.63 | 2738.06 |
| 50,000 | 38.61 | 57.38 | 88.83 | 383.27 | 756.06 | 3765.36 | 7380.50 |
| 100,000 | 130.05 | 205.89 | 335.81 | 1499.39 | 2977.47 | 14880.21 | 29145.10 |
| 300,000 | 1109.28 | 1828.38 | 2955.37 | 13329.59 | 26521.93 | - | - |
| 500,000 | - | - | 8152.81 | 36689.09 | - | - | - |
| 1,000,000 | - | - | 32499.53 | 146168.41 | - | - | - |

Supplementary Table 5: Wall-clock runtimes in seconds of *HybridGWAIS* logistic regression test with a different number of variants ($M$) and a different number of samples ($N$) using combined FPGA and GPU acceleration with an Alpha Data ADM-PCIE-8K5 PCIe FPGA accelerator board (containing a Xilinx Kintex UltraScale KU115 FPGA) and an Nvidia Tesla P100 GPU on our benchmark system.

| | $N$ = 1000 | 5000 | 10,000 | 50,000 | 100,000 |
|---|---|---|---|---|---|
| $M$ = 10,000 | 8.84 | 7.98 | 7.45 | 9.65 | 14.11 |
| 30,000 | 16.37 | 14.91 | 15.42 | 32.96 | 60.02 |
| 50,000 | 32.74 | 30.54 | 30.11 | 77.39 | 146.11 |
| 100,000 | 110.83 | 101.99 | 100.77 | 275.48 | 536.02 |
| 300,000 | 774.58 | 846.60 | 861.49 | 2319.71 | 4565.70 |
| 500,000 | 2468.55 | 2340.93 | 2389.54 | 6368.25 | 12591.84 |
| 1,000,000 | 9425.43 | 8597.85 | 9194.76 | 25308.44 | 50108.28 |
| 2,000,000 | 37078.51 | 33638.87 | 34288.09 | 100783.12 | 199877.05 |

Supplementary Table 6: Wall-clock runtimes in seconds of *HybridGWAIS* logistic regression test with a different number of variants ($M$) and a different number of samples ($N$) using combined FPGA and GPU acceleration with two Alpha Data ADM-PCIE-8K5 PCIe FPGA accelerator boards (containing a Xilinx Kintex UltraScale KU115 FPGA each) and an Nvidia Tesla P100 GPU on our benchmark system.

| | $N$ = 1000 | 5000 | 10,000 | 50,000 | 100,000 |
|---|---|---|---|---|---|
| $M$ = 10,000 | 7.99 | 7.79 | 7.86 | 9.51 | 12.16 |
| 30,000 | 15.42 | 14.84 | 15.41 | 22.44 | 38.91 |
| 50,000 | 34.65 | 31.75 | 31.79 | 45.60 | 84.80 |
| 100,000 | 88.58 | 104.79 | 101.96 | 150.04 | 288.01 |
| 300,000 | 725.87 | 805.68 | 863.05 | 1188.80 | 2344.75 |
| 500,000 | 2151.84 | 1871.66 | 2294.49 | 3227.95 | 7074.00 |
| 1,000,000 | 6957.15 | 7602.57 | 7575.77 | 14148.78 | 25734.47 |
| 2,000,000 | 31138.10 | 37226.14 | 35574.14 | 51941.97 | 101012.59 |

Supplementary Table 7: Wall-clock runtimes in seconds of *HybridGWAIS* for multiple testing methods and combinations of multiple testing methods. All runs were executed on the same dataset with $M$ = 100,000 variants and $N$ = 100,000 samples using different acceleration methods (GPU-only with an Nvidia Tesla P100 GPU and combined FPGA and GPU acceleration with one or two Alpha Data ADM-PCIE-8K5 PCIe FPGA accelerator boards respectively (containing a Xilinx Kintex UltraScale KU115 FPGA each) and the Tesla P100 GPU) on our benchmark system. The deviation is measured in percent (%) from the runtime of the logistic regression test.

| Method | GPU accel. | | 1x FPGA + GPU accel. | | 2x FPGA + GPU accel. | |
|---|---|---|---|---|---|---|
| | runtime | dev. | runtime | dev. | runtime | dev. |
| Logistic Regression (LogReg) | 2965.62 | 0.00% | 532.32 | 0.00% | 286.30 | 0.00% |
| BOOST | 2960.42 | -0.18% | 531.64 | -0.13% | 285.83 | -0.16% |
| Linkage Disequilibrium (LD) | 2951.15 | -0.49% | 531.33 | -0.19% | 285.46 | -0.29% |
| Mutual Information (MI) | 2948.55 | -0.58% | 531.03 | -0.24% | 285.08 | -0.43% |
| Information Gain (IG) | 2950.01 | -0.53% | 530.81 | -0.28% | 285.23 | -0.37% |
| LogReg + LD | 2974.76 | 0.31% | 533.07 | 0.14% | 287.44 | 0.40% |
| LogReg + BOOST | 2984.44 | 0.63% | 532.98 | 0.12% | 287.45 | 0.40% |
| LogReg + BOOST + LD | 2994.69 | 0.98% | 533.73 | 0.26% | 287.87 | 0.55% |
| LogReg + BOOST + LG + MI + IG | 3008.22 | 1.44% | 534.76 | 0.46% | 288.89 | 0.90% |

### References

[1] Apache HTTP Server Project, https://httpd.apache.org.

[2] PostgreSQL: The World's Most Advanced Open Source Relational Database, https://www.postgresql.org.

[3] Kässens, J. C. Qmanager, https://github.com/ikmb/qmanager.git.

[4] Wienbrandt, L. HybridGWAIS, `https://github.com/ikmb/hybridgwais.git`.

[5] A simple PHP WebAuthn (FIDO2) server library, https://github.com/lbuchs/WebAuthn.

[6] Purcell, S., Neale, B., Todd-Brown, K., et al. (September, 2007) PLINK: A Tool Set for Whole-Genome Association and Population-Based Linkage Analyses. *American Journal of Human Genetics,* **81**, 559–575.

[7] Wan, X., Yang, C., Yang, Q., et al. (2010) BOOST: A Fast Approach to Detecting Gene-Gene Interactions in Genome-wide Case-Control Studies. *Am. J. Hum. Genet.,* **87**(3), 325–340.

[8] Jakulin, A. and Bratko, I. (2003) Analyzing Attribute Dependencies. *Knowledge Discovery in Databases: PKDD 2003,* pp. 229–240.

[9] Jakulin, A. and Bratko, I. Quantifying and Visualizing Attribute Interactions: An Approach Based on Entropy. (2004).

[10] Wienbrandt, L., Kässens, J. C., Hübenthal, M., and Ellinghaus, D. (2019) 1000x Faster than PLINK: Combined FPGA and GPU Accelerators for Logistic Regression-based Detection of Epistasis. *Journal of Computational Science,* **30**, 183–193.

[11] Wienbrandt, L., Kässens, J. C., et al. (2017) Fast Genome-Wide Third-order SNP Interaction Tests with Information Gain on a Low-cost Heterogeneous Parallel FPGA-GPU Computing Architecture. *Proc. Computer Science,* **108**, 596–605.
